# Inferring Local Protein Structural Similarity from Sequence Alone

**DOI:** 10.1101/2025.11.24.690129

**Authors:** Zinnia Ma, Javier Espinoza Herrera, Neville P. Bethel, Adrian Jinich

**Affiliations:** University of California, San Diego

## Abstract

Detecting structural similarity at the local level between proteins is central to understanding function and evolution, yet most approaches require 3D models. In this work, we show that protein language models (pLMs), solely using sequence data as input, implicitly capture fine-grained structural signals that can be leveraged to identify such similarities. By mean-pooling residue embeddings over sliding windows and comparing them across proteins with cosine similarity, we find diagonal patterns that reflect locally aligned regions even without sequence identity. Building on this insight, we introduce a framework for detecting locally aligned structural regions directly from sequences, supporting the development of scalable methods for structural annotation and comparison.

## 1 Introduction

Understanding protein function and evolution often hinges on structural similarity, as proteins with comparable folds often perform related biochemical activities, even in the absence of sequence similarity [1]. Detecting such similarities is also critical in practice, supporting annotation of uncharacterized proteins, rational engineering, and drug discovery [2–5]. Traditional methods for identifying structural similarity, such as TM-align and DALI (Distance Matrix Alignment), align three-dimensional protein models obtained from experimental determination or computational prediction [6–8]. More recent tools, such as Foldseek, generate compact representations of protein structures to enable rapid comparisons; however, they still require a three-dimensional structural model as input [9].

In contrast, methods that solely rely on sequence have emerged; however, these are generally limited to detecting global fold-level similarities and remain insufficient at capturing local structural relationships [10–12]. More recently, pLMs such as ESM and ProtTrans have overcome this limitation by encoding structural and functional properties directly from sequence [13–15]. Despite being trained solely on large protein sequence corpora, their embeddings have been successfully applied to diverse downstream tasks, including secondary structure prediction, remote homology detection, contact prediction, and protein function annotation [16–19]. Thus, the ability to recover local structural similarity directly from sequence offers a practical advantage that bypasses the need for structural models. This motivates the central question of this work: whether pLM embeddings contain sufficient information to identify local structural similarity between proteins.

To address this question, we develop a sequence-only framework that detects locally similar structural regions using pLM-derived embedding. Our approach computes sliding window embeddings for two proteins, constructs a window-similarity matrix, enhances the resulting signal using a sigmoid-based transformation, and then identifies high-scoring local regions using a Smith-Waterman-style alignment procedure. This enables us to recover segments that are similar in structure, even when the full-length sequences differ substantially. We test a series of pLMs, including ProtT5, ESM-2 (3B) and (15B), CARP, and ProstT5 [12, 13, 20, 21]. Among them, ProtT5 and ProstT5 provided the clearest local structural signals. Applied to the MALISAM benchmark of structural analogs (protein pairs that share similar local folds despite lacking common ancestry) [22], our approach identifies local regions with high structural similarity, achieving TM-scores above 0.5 in many cases despite low global structural similarity. Comparative experiments further show that our method outperforms baseline methods based on residue alignment and predicted 3Di sequences, demonstrating that local structural information is well encoded in sequence-only ProstT5 embeddings. Taken together, our results establish a scalable, sequence-only framework for detecting local structural analogy that offers performance comparable to methods requiring explicit structural models, while providing a lightweight and complementary alternative.

## 2 Related Work

Foldseek represents the most efficient and widely used method for large-scale structural alignment. This method encodes protein structures as 3Di (3D interaction) sequences, which represent each residue by a structural state letter based on the 3D structure, facilitating high-throughput and accurate comparisons [9]. Recent sequence-based approaches such as ProstT5 and ESM-2 3B 3Di aim to bypass the need for structural data by directly translating amino acid sequences into predicted 3Di representations, thereby leveraging Foldseek’s powerful alignment algorithm [11, 12]. However, due to the limited prediction accuracy of 3Di tokenization, currently around 60%, these methods are better suited for global fold-level retrieval tasks than fine-grained fragment-level structural searches.

DeepBLAST is another existing model that could structurally align proteins using only sequence information [23]. This method is capable of aligning structurally homologous domains with low sequence identity, such as duplicated Annexin domains. However, because it is based on the Needleman–Wunsch algorithm, which is inherently designed for global alignment, DeepBLAST faces intrinsic limitations when homologous regions are restricted to small local segments. In such cases, enforcing a global alignment may dilute the signal of local structural similarity, making it harder to detect.

## 3 Methods

### Capturing the alignment signal via pairwise cosine similarity of sliding window embeddings

Given a protein sequence of length *n*, we first obtain per-residue embeddings of shape *n × d* using a pLM. To extract local contextual representations, we apply a sliding window of size *w* across the sequence, yielding in *n − w* + 1 overlapping segments. For each window, we perform mean pooling over its corresponding *w × d* local residue embeddings to produce a single *d*-dim window-level embedding. For a pair of proteins, we then compute cosine similarity between all pairs of window embeddings, generating a matrix where each entry reflects the similarity between specific local regions. This matrix serves as the basis for downstream analysis of structurally analogous regions.

### Enhancing alignment signal with sigmoid-based transformation

To convert the cosine similarity matrix into a reward matrix with enhanced contrast between aligned and misaligned regions, we applied a non-linear transformation function to emphasize high-similarity regions while penalizing low-similarity ones. Specifically, we used a scaled sigmoid function:

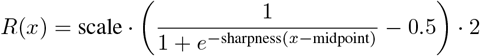

where *x* ∈ [ − 1, 1] is the normalized cosine similarity. Before applying the transformation, the raw similarity matrix was linearly rescaled to this range. The parameters were chosen to balance both signal sensitivity and robustness to noise.

### Detecting the alignment regions based on the Smith-Waterman algorithm

The Smith-Waterman algorithm is a dynamic programming method for local sequence alignment. In our approach, we adapt a Smith-Waterman-style algorithm to identify structurally aligned regions between a pair of proteins [24]. Specifically, we use the reward matrix obtained by applying the transformation described above to the raw cosine similarity matrix as the scoring basis in the Smith-Waterman framework (Figure 1). In this way, window pairs with high cosine similarity receive high match rewards, while dissimilar pairs are assigned strong mismatch penalties. To balance insertions and deletions, we introduce an additional indel penalty term. By completing the scoring matrix and performing traceback, we extract the highest-scoring aligned windows and map them back to the corresponding protein regions.

**Figure 1:**
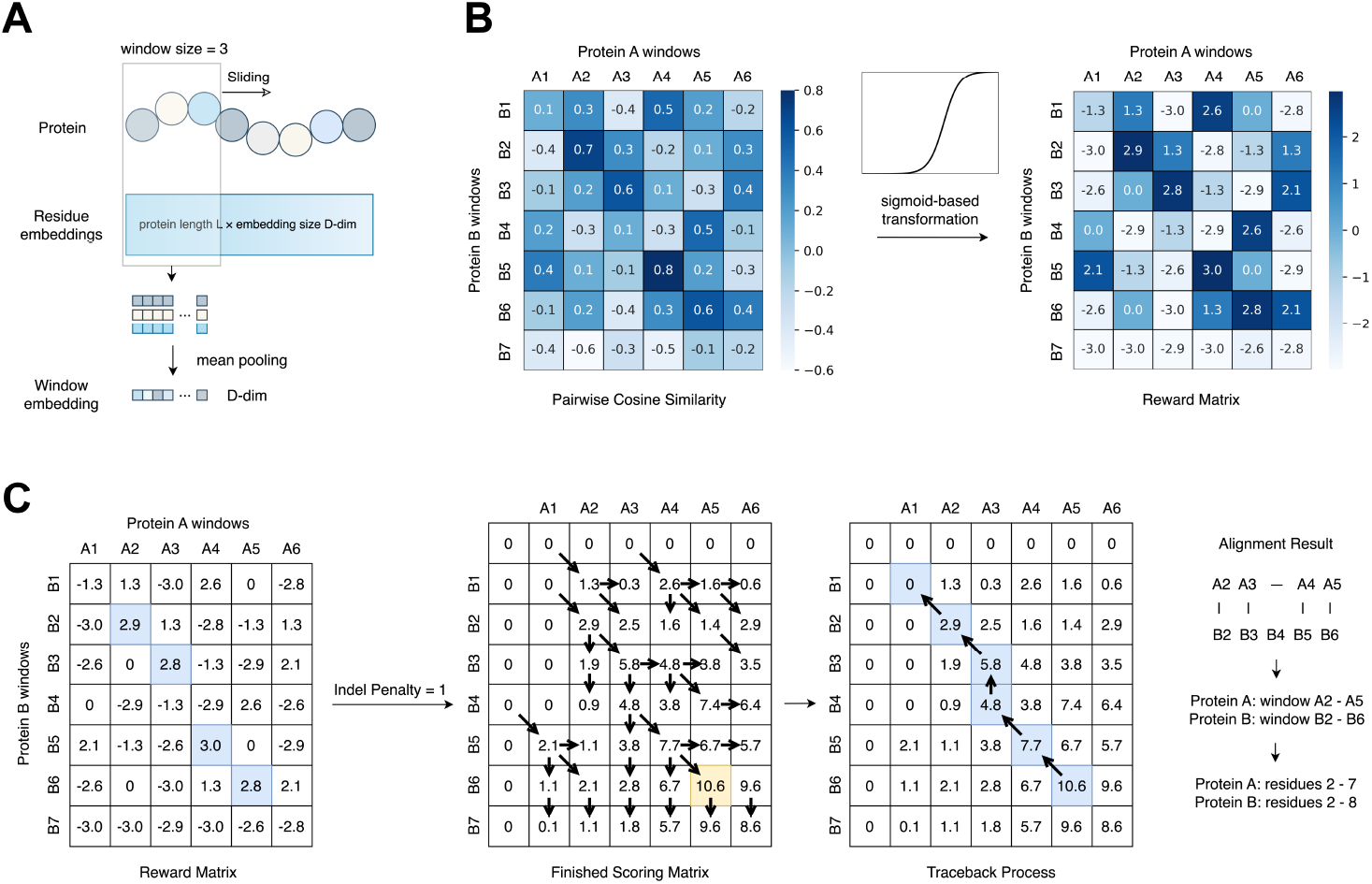
Workflow of the alignment method. (A) Extract window embeddings from pLM-derived residue embeddings. (B) Sigmoid-based transformation for signal enhancement. (C) Alignment based on the Smith-Waterman algorithm using a predefined reward matrixx. Cosine similarities between sliding window embeddings are transformed into a reward matrix, which is then used by a Smith-Waterman-style algorithm to identify structurally aligned regions. The figure demonstrates how our method reveals alignment signals embedded in the original pairwise cosine similarity matrix. The traceback path and corresponding alignment signal are highlighted in blue, while the maximum score in the scoring matrix is marked in yellow. The pseudocode of the algorithm is provided in Appendix 1.

## 4 Experiments

We evaluated our approach with the MALISAM database, a curated benchmark of 130 protein pairs from experimentally determined structures [22]. Unlike homologous proteins, which share features through common ancestry, MALISAM emphasizes analogous proteins that evolved similar folds independently. Due to their lack of shared ancestry, these protein pairs exhibit significantly lower sequence identity compared to structural homologs. Each pair consists of two sequence regions that align structurally even though their corresponding PDB structures differ at the global fold level. Each aligned region was defined using a combination of manual inspection and computational annotation. To curate the dataset, hybrid and core motifs were first identified across SCOP domains and then used as queries in DALI searches against a culled PDB set restricted to <50% sequence identity, ensuring that the resulting analog pairs reflect structural rather than evolutionary similarity. These motifs provide a rigorous standard for testing whether sequence-based embeddings can recover structural similarity.

### 4.1 Heatmap of embedding cosine similarity reveals structurally analogous regions

To test whether pLM embeddings capture structural similarity signals in the absence of high sequence identity, we generated pairwise cosine similarity matrices by comparing sliding windows across two analogous protein sequences. Each window corresponds to the average per-residue embedding from ProtT5 over five amino acids. In these matrices, higher cosine similarity values appear as continuous diagonals, suggesting alignment-like relationships between protein regions. Notably, these patterns emerge even when the sequences lack detectable homology, as with protein pairs in the MALISAM dataset containing structural analogs.

The two representative examples shown in Figure 2 illustrate these observations. For the pair 1a2z–1ghh in panel A, the diagonals in the similarity matrix align with *α*-helices in the 3D structures, showing that embedding-derived similarities directly coincide with secondary structure features. In contrast, the pair 1slc–1sq9 in panel B highlights how the predominance of *β*-sheets gives rise to multiple diagonals, again capturing structural alignment despite sequence divergence. Together, these cases demonstrate that windowed averages of protein embeddings effectively reflect three-dimensional organization, reinforcing their value in identifying structurally analogous regions in proteins with little to no sequence similarity.

**Figure 2:**
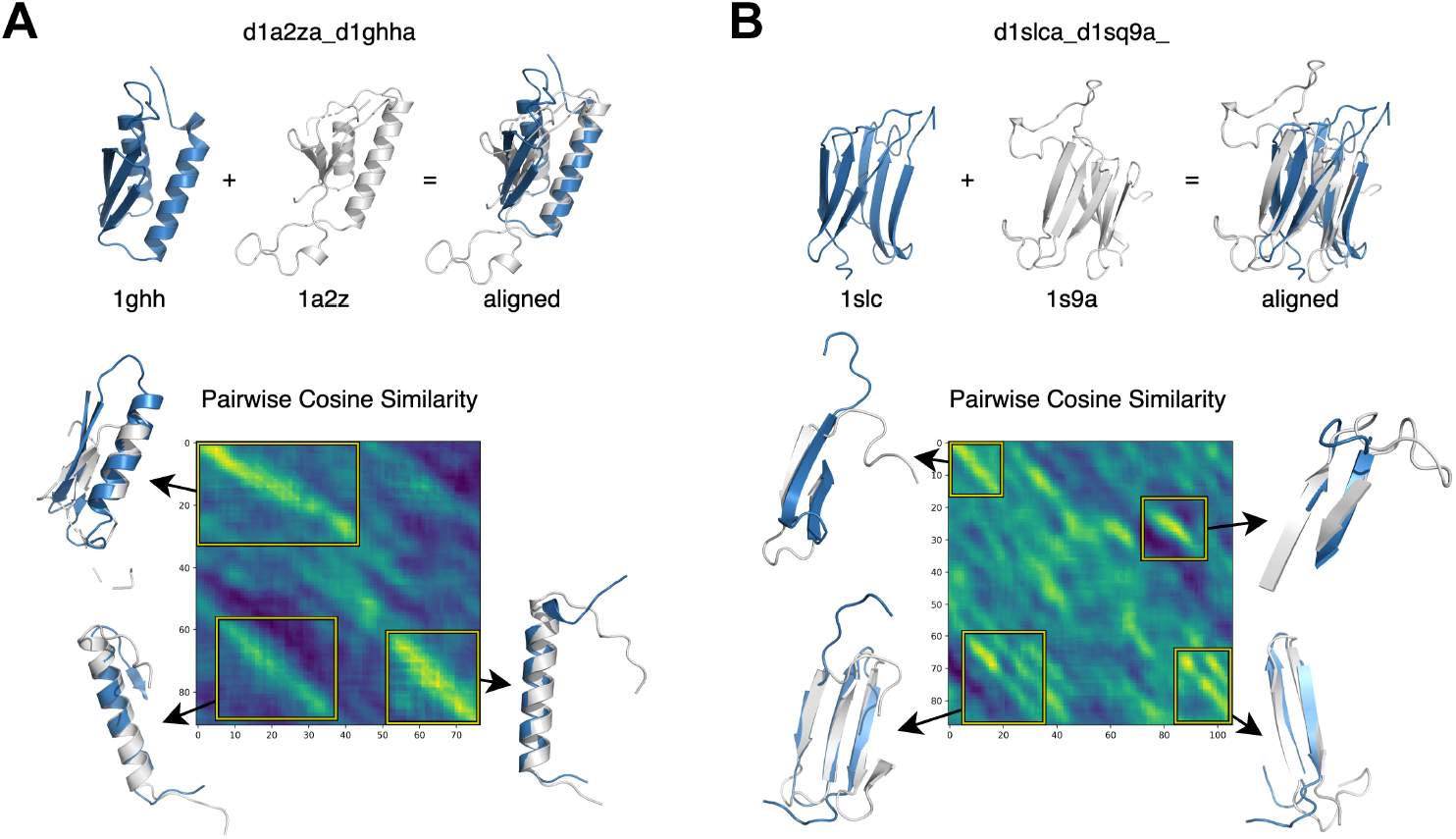
Heatmaps of cosine similarity across protein sequence pairs calculated using a sliding window of 5 residues. The y-axis represents the starting position of each window in the first protein, while the x-axis represents the starting position in the second protein. The heatmaps highlight structural correspondence, exemplified by (A) *α*-helices and (B) *β*-sheets in different protein pairs.

### 4.2 Comparing the performance of different pLMs in capturing fine-grained structural information

We continued by comparing the performance of different pLMs in capturing structural similarity signals. For each model, we compute pairwise cosine similarity matrices for all aligned protein pairs in the MALISAM dataset using a sliding window approach with a window size of 5. The resulting matrices are visualized as heatmaps. Representative examples are shown in Figure 3 and Appendix B.1. This qualitative comparison enables us to visually assess the strength and clarity of the structural signals captured by different pLMs. To more reliably assess the correspondence between the captured signals and the ground truth alignments, we further perform a quantitative evaluation. Specifically, we compute precision, recall, and F1-score by comparing the observed signal patterns in the heatmaps to the ground truth structural alignments. The detailed results are presented in Appendix B.2.

**Figure 3:**
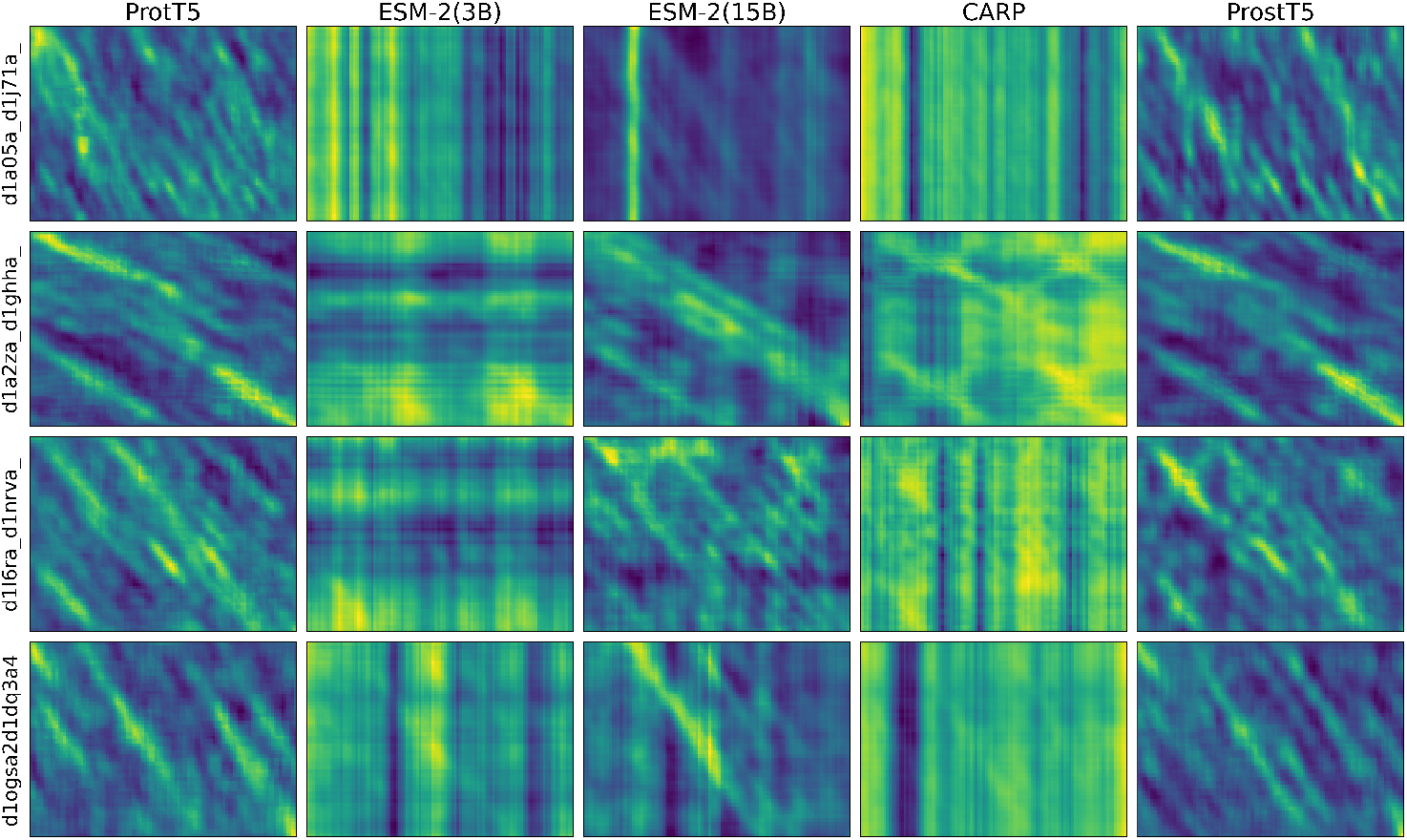
Comparison of pairwise cosine similarity patterns across different pLMs. Each column corresponds to a pLM: ProtT5, ESM-2 (3B), ESM-2 (15B), CARP, ProstT5, from left to right. Each row represents a randomly selected aligned protein pair from the MALISAM dataset, with the pair name shown on the left. The matrices are computed using a sliding window of size 5, and visualized to assess the reliability and clarity of alignment signals produced by different models.

The pLMs evaluated are ProtT5, ESM-2 (3B), ESM-2 (15B), CARP, and ProstT5. Among them, ProtT5 refers specifically to ProtT5-XL-U50 from the ProtTrans series, which adopts a T5-style architecture and contains approximately 3B parameters. The two ESM-2 variants utilize a BERT-style architecture with 3B and 15B parameters, respectively. CARP, in contrast, employs a ByteNet-style architecture and is significantly smaller, with only 640M parameters. ProstT5 differs from the other models in that it incorporates structural information during training. However, this structural signal is encoded in the form of 3Di sequences, allowing the model to remain purely sequence-based. Importantly, obtaining embeddings from ProstT5 still requires only the raw protein sequence as input. ProstT5 also adopts a T5-style architecture and is obtained by fine-tuning ProtT5-XL-U50.

From Figure 3, we observe that ProtT5 and ProstT5 consistently produce clear and stable diagonal signals, indicating well-aligned and coherent structural correspondence across the majority of protein pairs. In contrast, ESM-2 (3B) and CARP often generate noisy, grid-like artifacts, failing to capture consistent structural similarity. ESM-2 (15B) demonstrates noticeable improvement over the 3B variant but still exhibits degraded performance in certain cases. For example, the vertical streaks in the 1a05-1j7a protein pair illustrate one such failure mode. These patterns are consistently observed in the more extensive set of comparisons provided in Appendix B.1. Taken together, these results suggest that, from a qualitative perspective, ProtT5 and ProstT5 are the most robust and reliable models for capturing structural patterns. This may be attributed to their T5-style architecture. Notably, although ESM-2 (3B) shares a similar parameter scale and is also trained with a masked language modeling objective, it performs substantially worse on this task, which suggests that architecture may be a key contributing factor.

Further supporting the qualitative findings, the quantitative results in Appendix B.2 show that ProtT5 and ProstT5 indeed provide a more favorable balance between precision and recall compared to the other models. However, between the two, ProstT5 performs significantly better, achieving noticeably higher precision and F1-score. This indicates that while both models produce heatmaps with clear structural signals, the signals from ProstT5 more accurately correspond to the ground truth aligned regions. Among the pLMs trained purely on sequence data, ProtT5 achieves the strongest performance. In contrast, ProstT5 benefits from the explicit incorporation of structural information during fine-tuning, which enables the model to better learn how to generate embeddings that capture alignment-relevant features. This structural supervision leads to a clear performance gain over ProtT5 on this task. Accordingly, in the subsequent experiments, we use ProstT5 as the pLM for generating residue embeddings.

### 4.3 Effect of applying the sliding window and the sigmoid-based transformation

To assess the effectiveness and necessity of using the sliding window and sigmoid-based transformation in our pipeline, Figure 4 illustrates how different window sizes and the transformation affect the cosine similarity matrices, using the protein pair 1g99–1gqy as an example. The four columns correspond to sliding window sizes of 1, 5, 10, and 15, respectively. The first row shows the raw cosine similarity matrices computed directly from the window embeddings, while the second row presents the transformed matrices, using the sigmoid-based transformation described in Section 3. As shown, the strongest signal in the raw matrices, which has a cosine similarity of only around 0.5, is significantly highlighted after transformation in the reward matrices. This enhancement makes the key regions more distinguishable from the surrounding noise. In contrast, noise-prone regions in the top-right and bottom-left corners are strongly suppressed, remaining dark blue to indicate high penalties.

**Figure 4:**
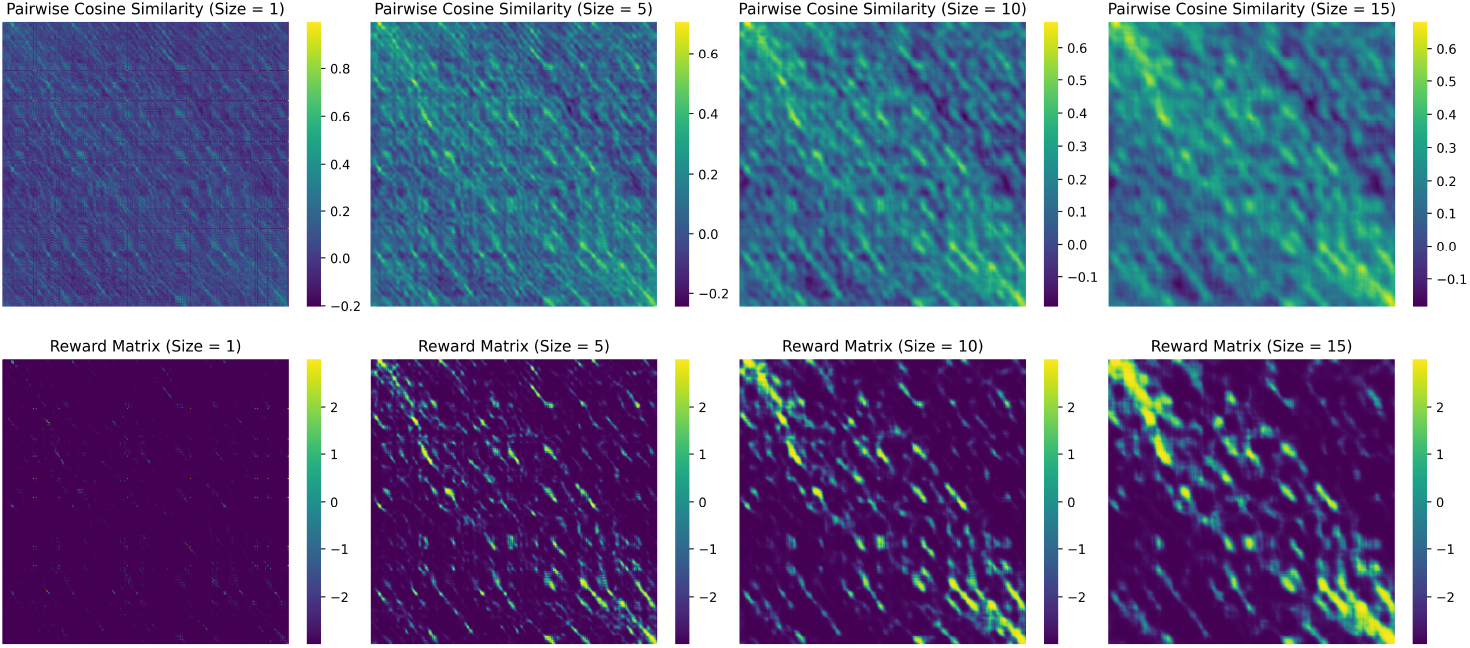
Comparison of cosine similarity matrices before and after sigmoid-based transformation across different sliding windows. The top row corresponds to the pairwise cosine similarity matrices before the transformation for protein pair 1g99-1gqy at different sliding window sizes. The bottom row shows the matrices after the transformation was performed using a sigmoid function.

Without using a sliding window as exemplified in the upper left of the figure (i.e., window size 1), the resulting matrix shows continuous signals that are equal across the grid. Thus, after the transformation, the resulting matrix is left with mostly penalized pairs, leaving little to no continuous signal. Using a sliding window amplifies the signal between sequences within a pair, making the aligned regions stand out from the background noise successfully. A trade-off exists between the size of the continuous signal and the fidelity, where the true signal is preserved more faithfully than with larger ones, where high cosine-similarity pairs dominate.

In the original raw cosine similarity matrix, many window pairs exhibit non-zero similarity scores, including a substantial amount of noise. The sigmoid-based transformation more precisely isolates truly informative signal regions. This transformation suppresses noisy, low-similarity scores by shifting them below zero, preventing the model from identifying overly large or spurious regions. At the same time, it accentuates high similarity scores, effectively highlighting salient regions in the reward matrix and enabling more accurate detection by subsequent algorithms.

### 4.4 Assessment of sliding window identified similarities proves detection of analogous structures

To evaluate the strength of the structural signal detected by our method, we first identified candidate aligned region pairs in the MALISAM dataset using our proposed workflow, followed by structural alignment to assess their validity. We employed ProstT5 as the pLM to extract embeddings, as our prior benchmarking demonstrated that ProstT5 most effectively encodes fine-grained structural information relevant to our task, outperforming other pLMs in this regard. Regarding hyperparameters, we used a sliding window of size 3. A sigmoid-based transformation was applied with a midpoint of 0.2, a sharpness of 9, and a scaling factor of 3. Details of the hyperparameter selection process are provided in Appendix B.3. For the alignment step, we adopted an indel penalty of 10, following standard practices in local alignment. For each protein pair in the MALISAM dataset, we applied our method to sequences derived from the complete PDB structures and identified the region pair yielding the highest alignment score under our framework. To assess whether the detected regions are indeed structurally similar, we computed TM-scores for both the full-length protein structures and the corresponding detected regions of each protein pair. These scores reflect global and local structural similarity, and their comparison is shown in Figure 5.

**Figure 5:**
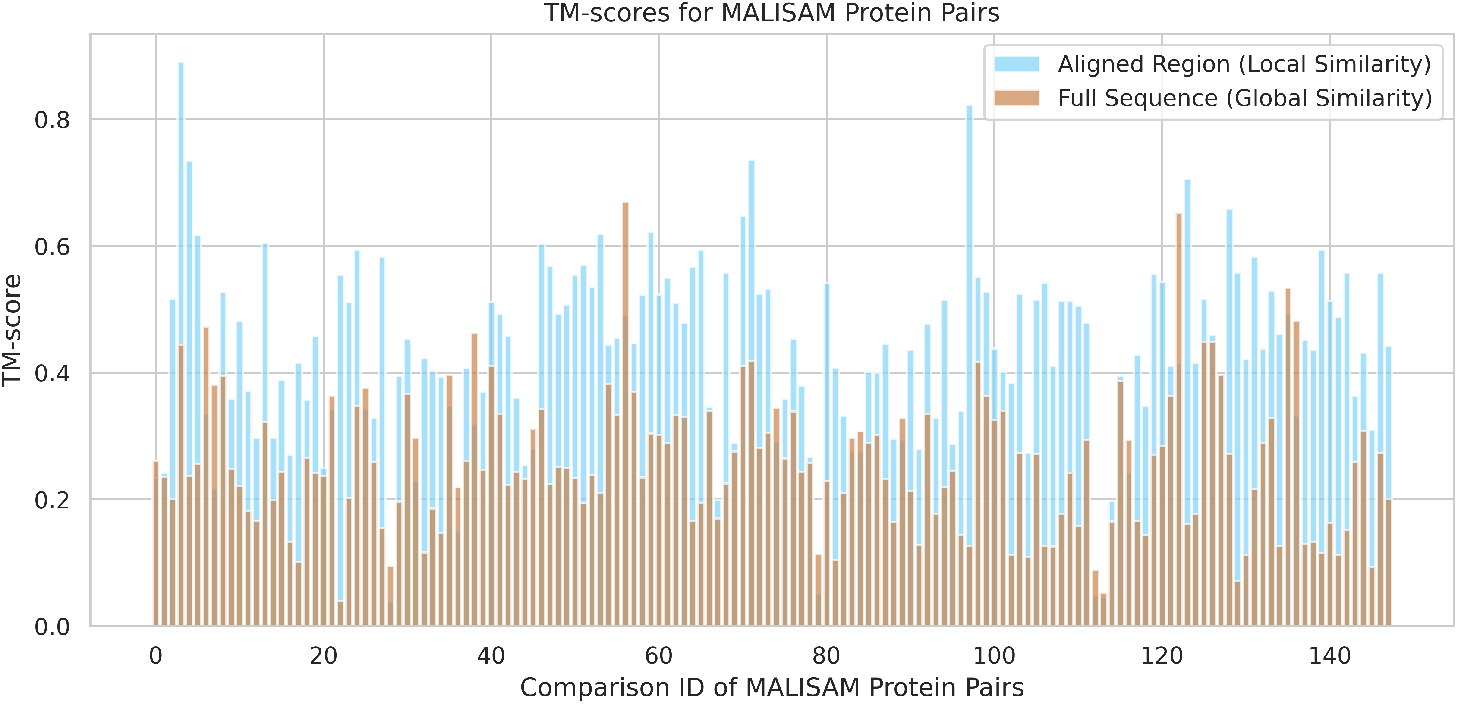
TM-score: Full-length structures vs. detected regions. Comparison of global and local structural similarity across all protein pairs in the MALISAM dataset.

**Figure 6:**
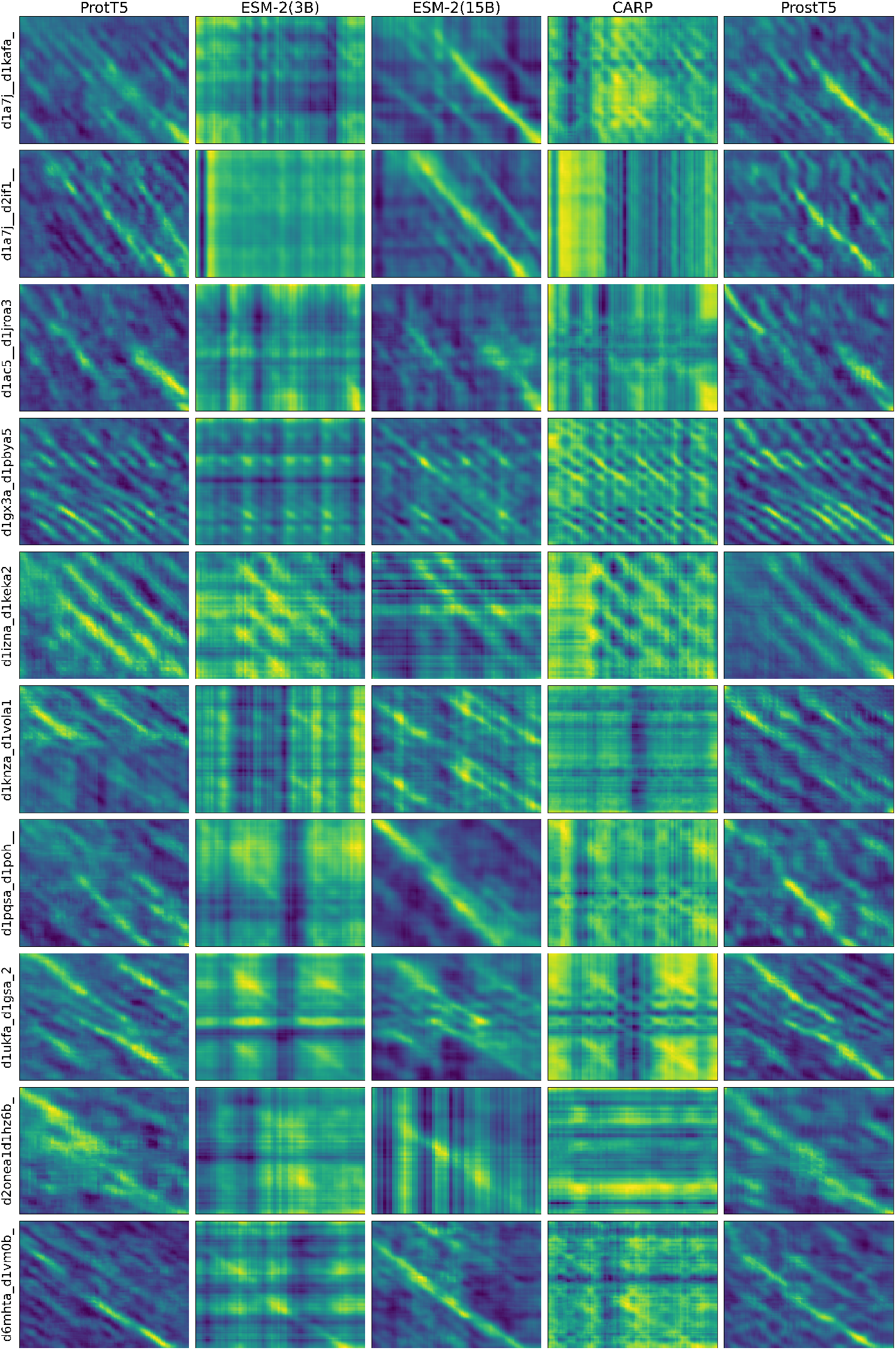
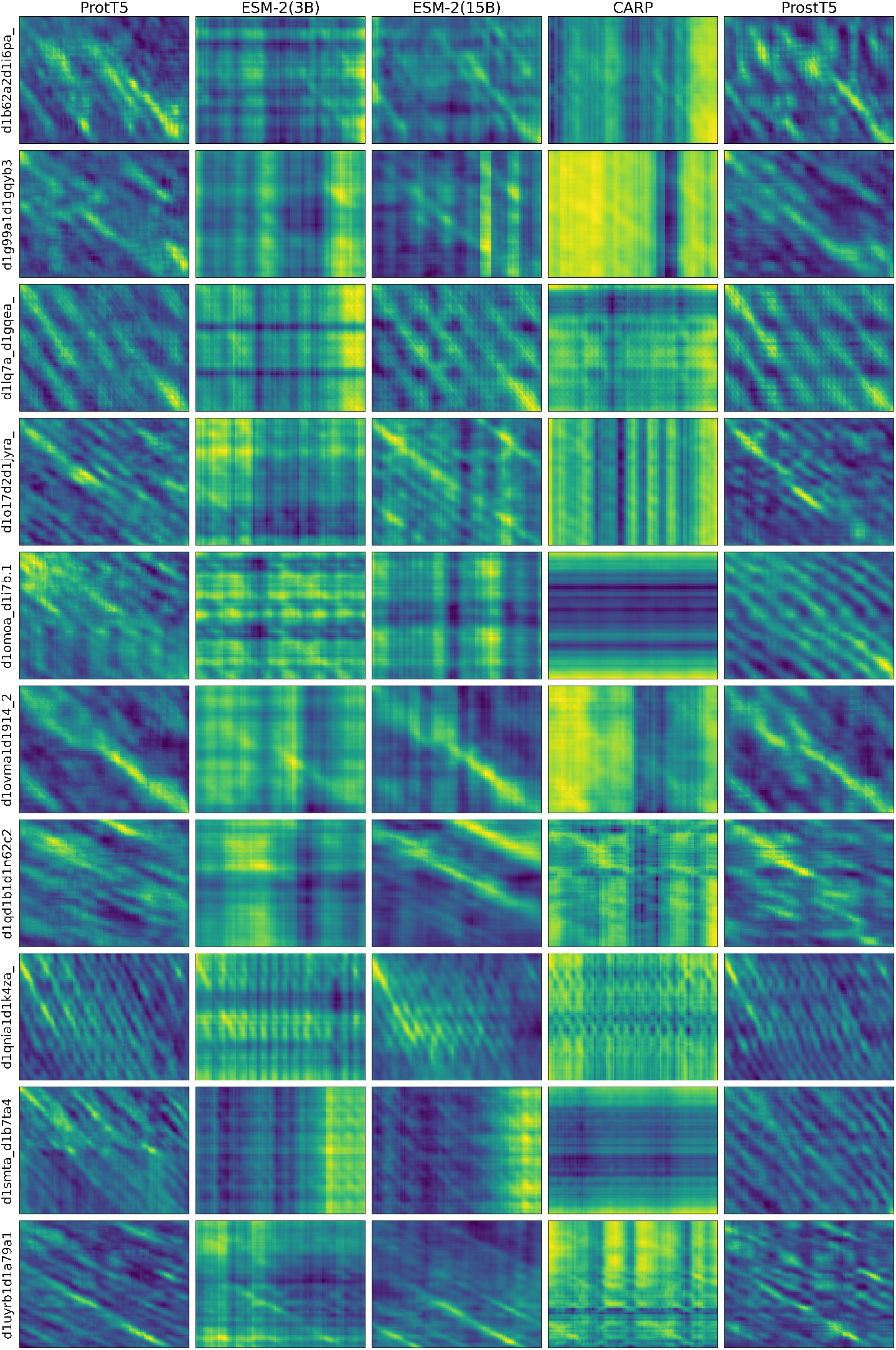
Extended comparison of pairwise cosine similarity patterns across different pLMs.

Among the 148 pairs of protein chains in the MALISAM dataset that contain aligned local structural regions, the global structural similarity, as measured by the TM-score between the full-length chains, remains low, averaging around 0.25. In contrast, the local similarity scores for the regions detected by our method are significantly higher. Specifically, 93 pairs achieve a TM-score above 0.4, with 53 of them exceeding 0.5. This indicates that our method is indeed capable of detecting aligned regions and motifs that are locally analogous in structure. However, there are also 4 pairs with local similarity scores below 0.1, and tuning the transformation hyperparameters did not lead to improved results for these cases. This suggests that the ability of ProstT5 to capture local structural similarity might be limited in some challenging cases, thereby influencing the downstream performance of our approach.

### 4.5 Comparison with residue and 3Di local alignment baseline methods

To evaluate the effectiveness of our workflow, we compared it against two representative baselines. The first baseline performs standard local alignment directly on raw protein sequences using Biopython’s PairwiseAligner, with the BLOSUM62 substitution matrix and gap penalties set to 11 (gap open) and 1 (gap extension), matching the default settings of blastp. The second baseline involves transforming the amino acid sequences into predicted 3Di sequences using ProstT5, followed by local alignment of the resulting 3Di representations using the same PairwiseAligner. For these 3Di alignments, we used Foldseek’s 3Di substitution matrix (mat3di.out) with gap penalties of 10 (gap open) and 1 (gap extension), matching the default gap costs used in Foldseek’s 3Di-based local alignment.

As shown in Table 1, we compare the average length of the predicted region pairs obtained by each method, as well as the number of pairs with TM-scores above or below specific thresholds. When comparing alignments based on amino acid sequences with those based on predicted 3Di sequences, the latter shows significantly better performance. Under comparable predicted length ranges, the method using predicted 3Di sequences produces nearly twice as many region pairs with TM-scores above 0.5 compared to the amino acid-based approach. Moreover, among methods that use the Prost T5 encoder, our sliding window + transformation + Smith-Waterman-like alignment approach outperforms the alternative that first decodes the embeddings into 3Di sequences and then aligns the resulting sequences. This suggests that the continuous embedding space captures local structural information more effectively than the discretized 3Di representation.

**Table 1:**
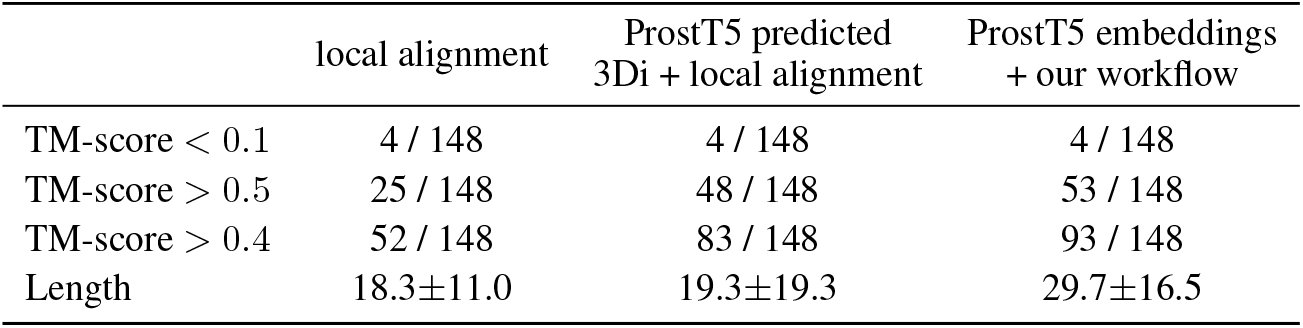
Comparison of methods based on TM-scores and lengths of detected local regions.

|  | local alignment | ProstT5 predicted<br>3Di + local alignment | ProstT5 embeddings<br>+ our workflow |
| --- | --- | --- | --- |
| TM-score < 0.1 | 4 / 148 | 4 / 148 | 4 / 148 |
| TM-score > 0.5 | 25 / 148 | 48 / 148 | 53 / 148 |
| TM-score > 0.4 | 52 / 148 | 83 / 148 | 93 / 148 |
| Length | 18.3±11.0 | 19.3±19.3 | 29.7±16.5 |

Our method can produce different results depending on the choice of hyperparameters. When the window size is fixed, the three hyperparameters related to the sigmoid-based transformation do not have a single clearly optimal setting. Instead, they primarily determine where the results fall along the tradeoff curve between region length and TM-score. To facilitate a fairer comparison with the other two baselines, we selected a hyperparameter configuration that results in relatively shorter region lengths: a midpoint of 0.2, a sharpness of 9, and a scaling factor of 3. This setting yields a mean region length of around 30 amino acids, which keeps the length reasonably close to those of the other two methods while still effectively capturing biologically meaningful motifs, about 10 amino acids longer than the regions identified by the other baselines. At the same time, our method continues to detect more high TM-score pairs, identifying 10 and 5 more pairs with TM-scores greater than 0.4 and 0.5, respectively, compared to the method based on predicted 3Di sequences.

Overall, these results validate the effectiveness of our method. They also indicate that information relevant to local structural similarity is already well encoded in the ProstT5 embeddings. While decoding the 3Di sequence can indeed make the approach more compatible with the Foldseek toolchain for structure similarity search, it is not necessarily the most effective way to exploit the structural information embedded in the representations.

## 5 Conclusion

In this work, we investigate the fine-grained structural information encoded in pLM embeddings and explore their potential for detecting local protein structural similarity. We present a scalable and zero-shot framework for detecting structurally aligned regions directly from sequence data, and provide initial validation of this approach using a rigorous benchmark, the MALISAM dataset, which comprises protein pairs with low sequence similarity. In many cases, the identified regions achieve TM-scores above 0.5, indicating strong structural correspondence.

Our method builds on the key observation that pairwise cosine similarity computed from pLM embeddings using a sliding window can capture structurally analogous regions. As illustrated in Section 4.1, this signal consistently emerges across protein pairs from different structural classes, suggesting that this methodology is generalizable across protein folds. Among the models evaluated, ProtT5 and ProstT5 yielded the most distinct and reliable alignment patterns, suggesting a strong inductive bias of the T5 architecture toward capturing local structural features.

When comparing our approach to baseline methods, we find that although both use the same ProstT5 encoder, our framework outperforms the approach that decodes embeddings into 3Di sequences followed by local alignment. This observation further supports the idea that information relevant to local protein structural similarity is largely preserved in the sequence-derived embeddings themselves. Converting embeddings into 3Di sequences is not the only way to utilize this information, and may not be the most effective. Our framework illustrates an alternative approach that makes more direct use of the structural signals encoded in pLM embeddings.

## Acknowledgements

This work is HHMI Hanna Gray Fellowship (NB: #17718 AJ: #16787), UC San Diego NIH FIRST Grant (NB, AJ: NCI 1UA54CA272220), and the Powell-Bundle Fellowship (ZM). We also thank the San Diego Supercomputer Center (SDSC) and UCSD Physics Computing Facility for computational resources.

## A Supplementary Methods

### A.1 Algorithmic Details

#### Algorithm 1

Local Alignment (Smith-Waterman) Using Predefined Reward Matrix

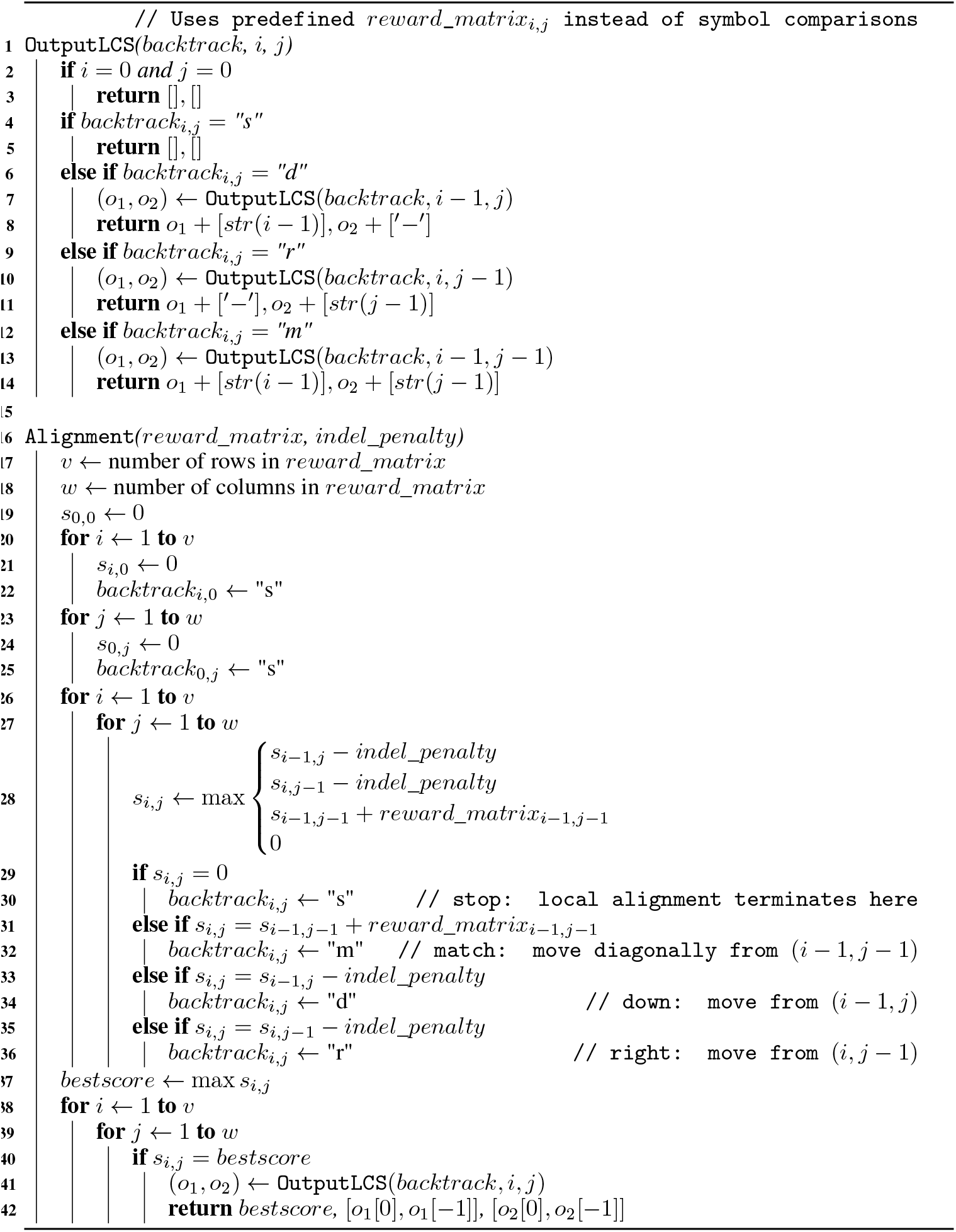

## B Supplementary Experiments

### B.1 Supplementary comparison of patterns across different pLMs

### B.2 Quantitative comparison of patterns across different pLMs

To quantify how well raw cosine similarity between embeddings can recover known structurally aligned regions, we used the TM-align–based alignments provided in the MALISAM entries as ground truth. For each protein pair, we parsed the corresponding file and generated an alignment matrix in which an entry is 1 only when both residues are aligned and belong to the analogous motif, and 0 otherwise. Independently, for each model, we computed an all-vs-all cosine similarity matrix between per-residue embeddings of the two sequences. Given a cosine similarity threshold, we then predicted residue pairs as aligned wherever the similarity matrix exceeded the threshold, and compared this predicted alignment to the ground-truth matrix to count true positives, false positives, and false negatives at the residue-pair level. From these counts, we computed precision, recall, and F1-score for each threshold, model, and protein pair.

Figure 7 summarizes the effect of cosine similarity thresholding on alignment precision, recall, and F1-score across the three models. Across all thresholds, the models exhibited the expected trade-off between precision and recall, though their performance profiles differed significantly. At low cosine similarity thresholds, recall remained maximal across models, reflecting the recovery of a large number of non-specific alignments. However, this resulted in very low precision. As the threshold increased, the precision of ProstT5, ProtT5, and ESM-2 (15B) improved modestly, peaking before declining to zero. In contrast, the precision of CARP and ESM-2 (3B) barely surpassed the baseline value and declined rapidly. Similarly, the latter two models failed to improve the F1-score beyond their initial value. These results are consistent with the visual patterns observed in the similarity matrices. ProstT5, ProtT5 and ESM-2 (15B) produced alignment-like diagonal regions, whereas CARP and ESM-2 (3B) produced diffuse straight-line patterns that did not suggest meaningful structural correspondence. Overall, these results suggest that ProstT5 offers the best balance between precision and recall, maintaining signal across a wider range of thresholds than ProtT5 and ESM-2 (15B). In contrast, CARP and ESM-2 (3B) are less effective at capturing local structural similarity from sequences alone.

**Figure 7:**
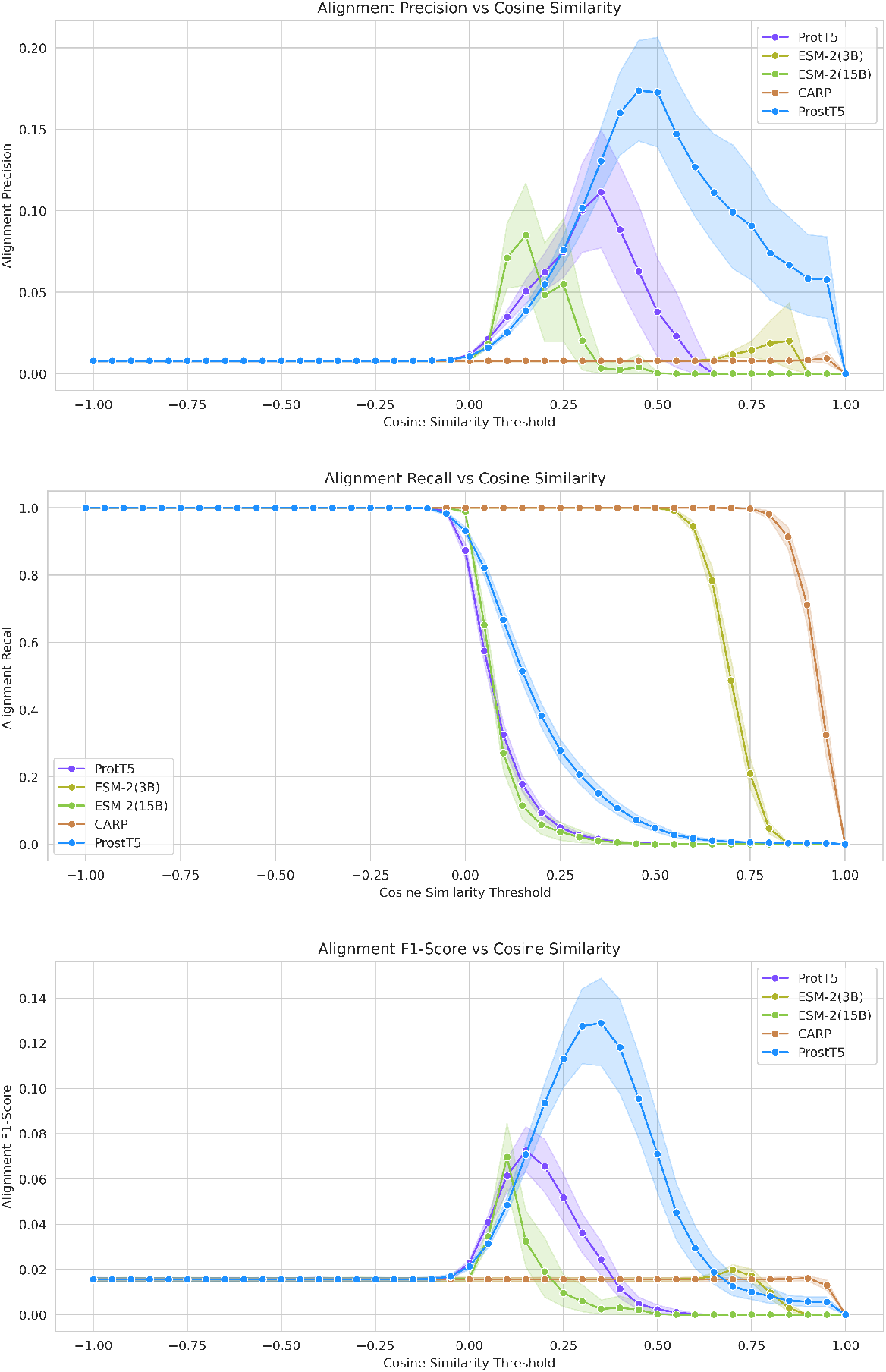
Pairwise alignment metrics for precision (top), recall (middle), and F1-score (bottom) across different cosine similarity thresholds for various pLMs.

### B.3 Hyperparameter selection

To identify the most suitable hyperparameters, we conducted an extensive comparison and search. For each window size, we performed a grid search over combinations of three hyperparameters used in the sigmoid-based transformation: midpoint, sharpness, and scale. Specifically, the midpoint was varied from 0 to 1 with a step size of 0.1, while both sharpness and scale were varied from 1 to 10 with a step size of 1, resulting in a total of 1100 combinations. For each combination, we applied our alignment method to all protein pairs, generating a pair of aligned regions for each, and subsequently computed the region length and TM-score for each pair.

Figure 8 presents results from hyperparameter search experiments. For each hyperparameter combination under each window size, we computed the average region length and TM-score across all protein pairs. Each point in the scatter plot corresponds to one such combination, with the color indicating the window size, as specified in the legend. We observe a consistent trend across the dots corresponding to each window size: higher TM-scores are generally associated with shorter region lengths. As the hyperparameters are adjusted to make the sigmoid function impose a stricter penalty, this leads to the detection of shorter regions. At the same time, such stricter filtering results in higher TM-scores, reflecting more confident alignments.

**Figure 8:**
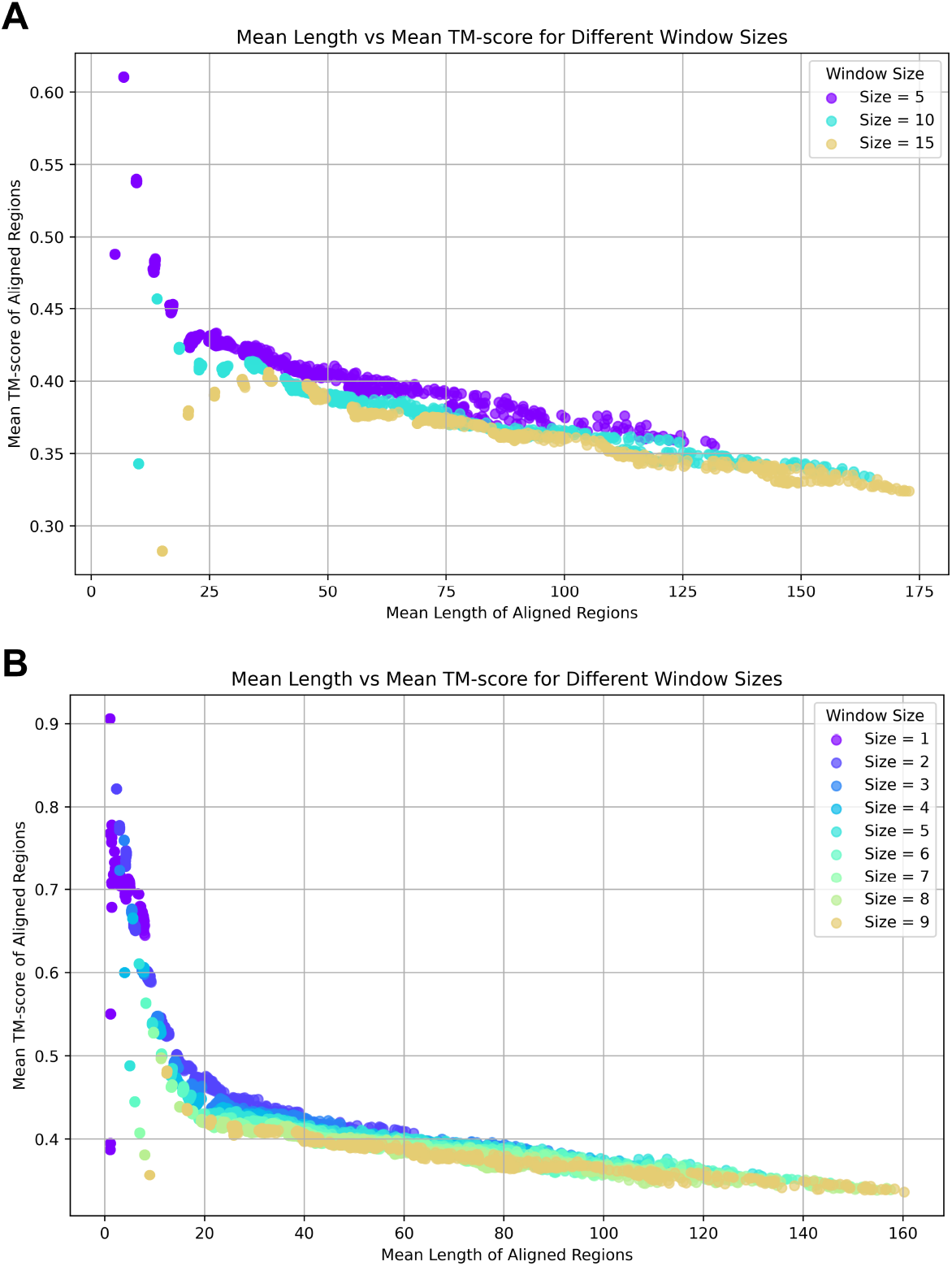
Grid search results for sigmoid-based transformation hyperparameters across different window sizes.

When comparing across different window sizes, as shown in Panel A, we find that the distributions of points shift noticeably: for a window size of 5, the points are concentrated more towards the upper-left region of the plot; for a window size of 15, they shift towards the lower-right; and window size of 10 lie in between. This observation is consistent with the results presented in Figure 4. Increasing the window size smooths the signals in the pairwise cosine similarity matrix, reducing the likelihood that small gaps disrupt contiguous aligned regions. Consequently, the detected regions tend to be longer, but this comes at the cost of lower TM-scores. Notably, for a fixed region length, points corresponding to window size 5 consistently achieve higher TM-scores, indicating better alignment quality. Based on this, we consider window size 5 to be a more desirable setting. To further refine our choice, we conducted a finer-grained search around this value, evaluating all window sizes from 1 to 9, as shown in Panel B.

The results reveal that the general trend still holds: smaller window sizes tend to yield points closer to the upper-left region of the plot, reflecting shorter but higher-quality alignments. However, some exceptions are observed. At window size 1, where no sliding window is applied, the lack of signal continuity prevents the detection of region pairs of sufficient length. Consequently, all hyperparameter combinations fail on most protein pairs. With window size 2, some valid regions begin to appear in a subset of cases, but no combination achieves consistent success across all pairs. It is only from window size 3 onward that certain hyperparameter settings yield successful region detections for the full dataset. Notably, starting at window size 5, all tested hyperparameter combinations are able to detect valid regions across all protein pairs.

When the window size is fixed, varying the combination of the three hyperparameters in the sigmoid-based transformation reveals a clear trade-off between detecting regions with higher TM-scores and identifying longer regions. A higher midpoint leads to more areas in the original pairwise cosine similarity matrix being suppressed, effectively classifying them as mismatches and penalizing them accordingly. This results in shorter detected regions. Increasing the sharpness amplifies already prominent signals, which encourages the traceback procedure to favor these stronger regions over other contiguous but weaker segments. A higher scale, on the other hand, amplifies the reward associated with longer regions, making it easier to overcome the fixed indel penalty (set to 10) and thereby reducing the likelihood of interruptions within the aligned region. Overall, there is no universally optimal combination of these hyperparameters, as the best choice depends on the desired balance between alignment accuracy and region length for different applications. In our experiments, we selected the hyperparameter setting that achieved the highest mean TM-score while still detecting valid regions for all protein pairs under a window size of 3. This led us to choose a midpoint of 0.2, a sharpness of 9, and a scaling factor of 3.

We further observed that the number of cases with TM-score greater than 0.5 or 0.4 is positively correlated with the average TM-score, while the number of cases with TM-score below 0.1 consistently remains limited to four specific cases. This suggests that when the pLM captures reasonably accurate underlying information for a given case, hyperparameter tuning can significantly influence the quality of the detected regions. In contrast, when the pLM fails to provide reliable underlying information, downstream tuning becomes largely ineffective. A closer examination of these four cases reveals that three of them involve protein pairs consisting of an artificial protein and a natural protein, suggesting that ProstT5 may still be limited in its ability to represent artificial proteins, which could explain the observed failure cases.

